# Experimental evolution to identify undescribed mechanisms of resistance to a novel cationic peptide antibiotic

**DOI:** 10.1101/2020.12.16.423161

**Authors:** A Santos-Lopez, MJ Fritz, JB Lombardo, AHP Burr, VA Heinrich, CW Marshall, VS Cooper

## Abstract

A key strategy for resolving the antibiotic resistance crisis is the development of new drugs with antimicrobial properties. The engineered cationic antimicrobial peptide WLBU2 (also known as PLG0206) is a promising broad-spectrum antimicrobial compound that has completed Phase I clinical studies. It has activity against Gram-negative and Gram-positive bacteria including infections associated with biofilm. No definitive mechanisms of resistance to WLBU2 have been identified. Here, we used experimental evolution under different levels of mutation supply and whole genome sequencing (WGS) to detect the genetic pathways and probable mechanisms of resistance to this peptide. We propagated populations of wild-type and mutator *Pseudomonas aeruginosa* in the presence of WLBU2 and performed WGS of evolved populations and clones. Populations that survived WLBU2 treatment acquired a minimum of two mutations, making the acquisition of resistance more difficult than for most antibiotics, which can be tolerated by mutation of a single target. Major targets of resistance to WLBU2 included the *orfN* and *pmrB* genes, previously described to confer resistance to other cationic peptides. More surprisingly, mutations that increase aggregation such as the *wsp* pathway were also selected despite the ability of WLBU2 to kill cells growing in a biofilm. The results show how the experimental evolution and WGS can identify genetic targets and actions of new antimicrobial compounds and predict pathways to resistance of new antibiotics in clinical practice.

## Introduction

The ability of a bacterial population to evolve resistance to an antibiotic depends on several factors including the availability of mutations that increase resistance and the strength of selection imposed by the compound. Experimental evolution coupled with whole genome sequencing (WGS) is a powerful strategy to characterize genetic mechanisms of resistance. Propagation of a bacterial population in the presence of an antibiotic will eventually select those clones that are capable of surviving the antibiotic exposure, and WGS of these populations or clones will reveal genetic causes of the resistance phenotype (Gullberg et al. 2011; Barbosa et al. 2019; Maltas and Wood 2019; Santos-Lopez et al. 2019). This method is especially powerful when studying cationic peptides, as multiple mutations may be needed to evolve resistance to them (Perron et al. 2006; Mehta et al. 2018; Spohn et al. 2019).

WLBU2 (also called PLG0206) is an engineered amphipathic alpha helix derived from the LL-37 peptide that inserts into the bacterial membrane and leads to cell death. WLBU2 is shown to be highly effective against ESKAPE (***<u>E</u>****nterococcus faecium*, ***<u>S</u>****taphylococcus aureus*, ***<u>K</u>****lebsiella pneumoniae*, ***<u>A</u>****cinetobacter baumannii*, ***<u>P</u>****seudomonas aeruginosa* and ***<u>E</u>****nterobacter spp*.) pathogens *in vitro* and *in vivo* (Paranjape et al. 2013; Deslouches et al. 2015; Lashua et al. 2016; Melvin et al. 2016; Mandell et al. 2017; Chen et al. 2018; Lin et al. 2018; Kumagai et al. 2019; Swedan et al. 2019; Di et al. 2020; Mandell et al. 2020). Antibiotic activity is typically measured against planktonic cells as biofilms are highly antibiotic tolerant, but WLBU2 has demonstrated a high activity against *P. aeruginosa* and *S. aureus* biofilms (Mandell et al. 2017; Chen et al. 2018; Mandell et al. 2020).

Despite challenging several Gram-negative and Gram-positive pathogens with WLBU2 *in vitro* and in animal models, only *P. aeruginosa* has been observed to develop resistance to WLBU2 after long periods of exposure (Deslouches et al. 2015). However, the mechanism of resistance remains undefined. Identifying the genes and the subsequent mutations that confer resistance to novel antibiotics is crucial as it enables more comprehensive understanding of the modes of action of this new antimicrobial compound and how resistance to them evolves. Further, these discoveries can increase our ability to predict the emergence of antimicrobial resistance (AMR) in clinical scenarios (Brockhurst et al. 2019; MacLean and San Millan 2019; MacLean 2020). We hypothesize that resistance to WLBU2 is rare because of its effectiveness and because it requires a specific combination of mutations. Therefore, a large mutation supply may be needed to generate the narrow range of combinations of resistance determinants. A large mutation supply could be generated by an increase in mutation rate, large population sizes, and/or long periods of time (van Dijk et al. 2017). Due to the effectiveness of WLBU2 at killing and keeping population sizes small, we anticipated that increased mutation rates or prolonged subinhibitory exposure would be the most likely means for the bacteria to generate a larger mutation supply. We tested each of these strategies using experimental evolution coupled with genomic sequencing to define mechanisms of resistance to this new cationic peptide.

## Results and Discussion

### Failure to evolve resistance using laboratory strains and standard protocols

To identify mechanisms of resistance to WLBU2, we propagated the laboratory strains of *A. baumannii* ATCC 17978 and *P. aeruginosa* PA14 that had no prior exposure to WLBU2. We followed a protocol successfully used previously with several pathogens and different antibiotics, in which large populations of bacteria are treated with increasing concentrations (Figure 1) (Hernando-Amado et al. 2019; Santos-Lopez et al. 2019; Trampari et al. 2019; Scribner et al. 2020; Santos-Lopez et al. 2021). We serially passaged both strains (bottleneck of ∼10^7^ cfu) in WBLU2 concentrations initially at half of the Minimum Inhibitory Concentration (MIC) and increasing two-fold every three days, up to four times the MIC. We propagated the same number of populations in the absence of WLBU2 to distinguish between adaptation to the environment and WLBU2-related mutations. Knowing that biofilm formation can influence the evolution of resistance (Ahmed et al. 2018; Santos-Lopez et al. 2019; Scribner et al. 2020), we implemented this regimen in both planktonic and biofilm cultures (see Figure S1, http://evolvingstem.org and (Santos-Lopez et al. 2019; Scribner et al. 2020) for details of the biofilm propagation). To test the general capability of the experimental design for selecting populations resistant to cationic peptides, we also propagated populations of *A. baumannii* in each lifestyle with increasing concentrations of polymyxin B. While all planktonic populations and four out of five biofilm populations survived in four times the MIC of polymyxin B, mainly through mutations in the *pmrABC* operon and in genes modifying lipid A (Table S1), no populations of *A. baumannii* nor *P. aeruginosa* survived exposure to the MIC of WLBU2. This suggested that there may be a narrow genetic pathway to gain WBLU2 resistance that could require more than one mutation.

**Figure S1.**
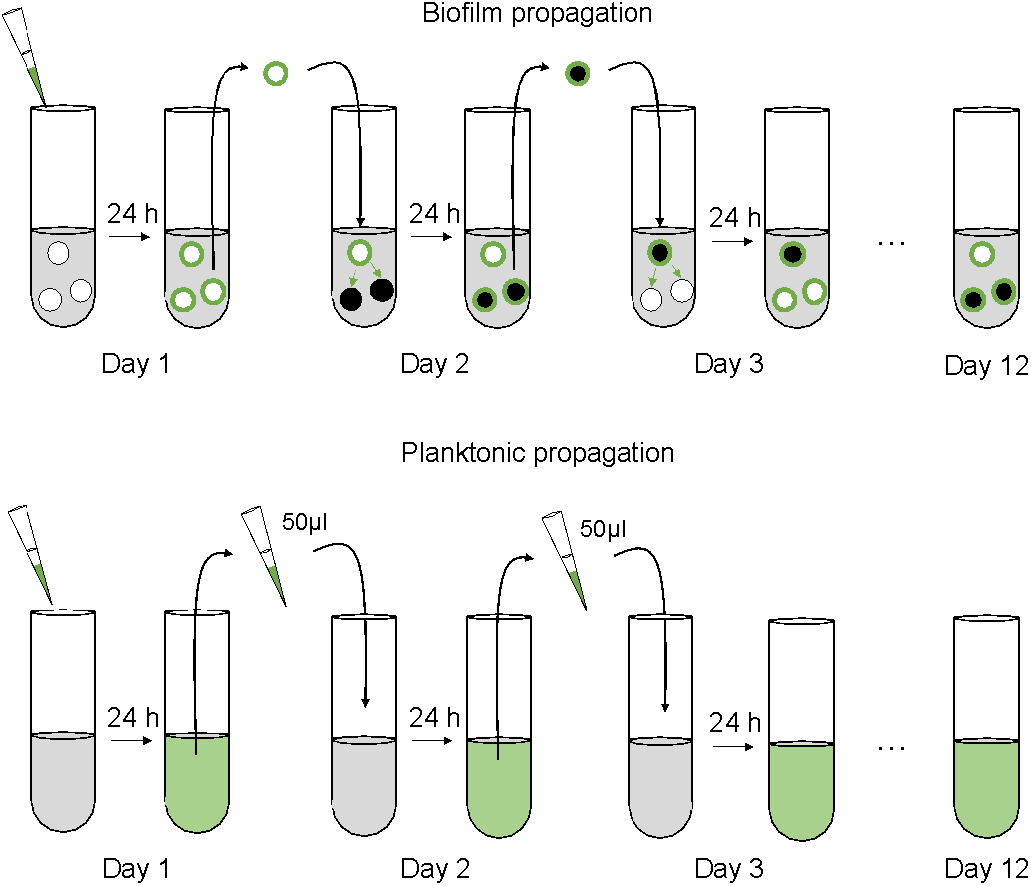
Schematic representation of the biofilm propagation (top) and the planktonic propagation (bottom) used to propagate the sensitive strains in presence of WLBU2. For biofilm populations (top), we transferred a polystyrene bead of the 24-hour culture to fresh media containing three sterile beads, which selects for adherent cells more than planktonic cells. Each day we alternated between black and white marked beads, ensuring that the bacteria were growing on the bead for 24 hr, which corresponds to approximately 6 to 7.5 generations/day. For the planktonic propagation (bottom), we serially passaged 50 µl into 5 ml of M9+ (dilution factor 100) which corresponds to approximately 6.64 generations per day.

**Figure 1:**
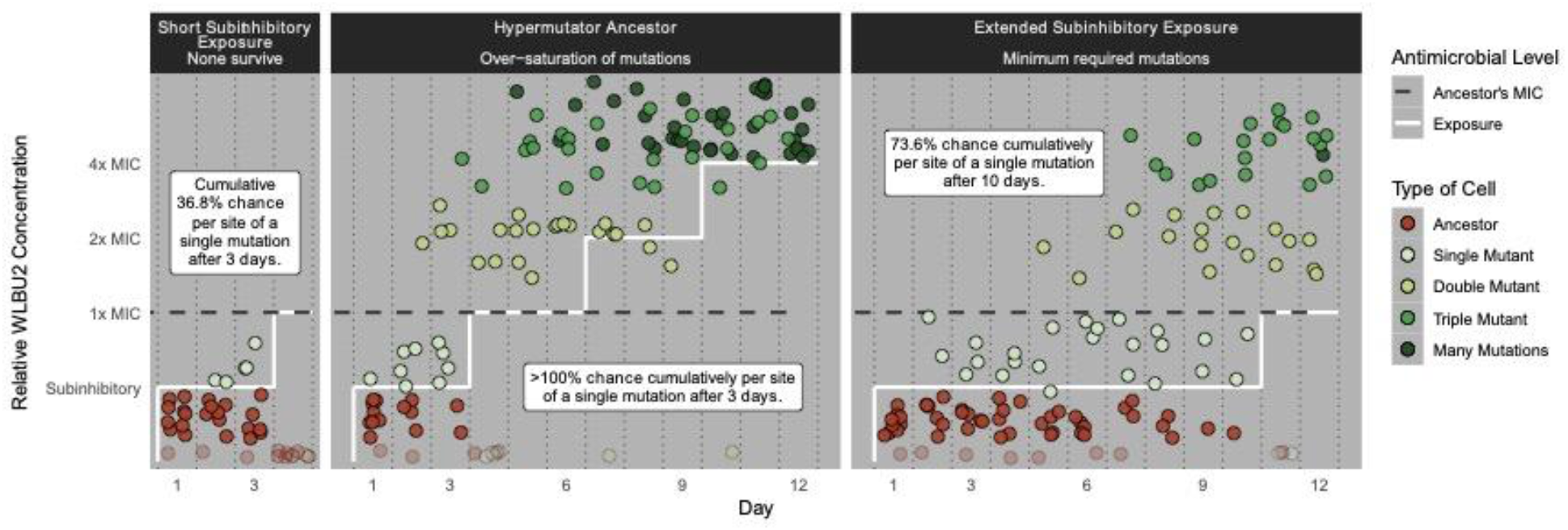
Schematic overview of the experimental setups. We exposed wild-type *P. aeruginosa* PA14 (WT) to three different WLBU2 treatments. In the first one (left panel), we grew populations in subinhibitory concentrations of WLBU2 for 3 days, and then we doubled this concentration to inhibitory concentrations that caused extinction. In the second experiment (middle panel), we used a hypermutator PA14 strain with a 100-fold greater mutation rate than WT. In the third one (right panel) we propagated WT for 10 days in subinhibitory concentrations of WLBU2, thus increasing the mutation supply by augmenting the number of generations. Dots simulate the expected number of mutants as calculated in Table 1.

The two main factors affecting the likelihood for a microbial population to evolve antibiotic resistance are (Hughes and Andersson 2017): i) the mutation supply that is determined by the product of the population size and mutation rate and ii) the strength of selection imposed by the antibiotic. If we assume a conservative mutation rate of 0.001 mutations per genome per generation, and a Poisson distribution of the mutations in the genome (Dillon et al. 2017), this experiment only had a cumulative probability of *∼*0.35 for *A. baumannii* and 0.29 for *P. aeruginosa* that any given nucleotide had been mutated at least once (in one cell) following three days of growth (∼36 generations) in subinhibitory concentrations, *i*.*e*. before being exposed to lethal concentrations (Tables 1 and S2 for detailed calculations). It is therefore possible that the correct mutation or combination of mutations needed to survive the antibiotic treatment had not occurred, or had been lost from the transfer bottleneck, before facing inhibitory concentrations. We therefore pursued two approaches to increase the number of mutations sampled in our *P. aeruginosa* populations to cultivate resistance to WLBU2: increasing the mutation rate and extending the duration of subinhibitory exposure (Figure 1).

**Table 1:** Mutation supply and distribution of the mutations in the three different experiments. We can calculate the mean mutations per nucleotide and the cumulative probability of a mutation per nucleotide knowing the mutation rate, the length of the chromosome, the number of divisions/days, and the length of exposure before facing inhibitory concentrations of antibiotic. ^a^ Calculated in (Harris et al. 2021).^b^ Days propagated before facing inhibitory concentrations of WLBU2. ^c^ Calculated as the sum of cell divisions to regrow the population each day following dilution into fresh culture. ^d^ Total cell divisions multiplied by mutation rate. ^e^ Total mutations divided by chromosome size. ^f^ Based on Poisson distribution of the mutations. For details of the calculations, see Table S2.

|  | Previously<br>used<br>protocol | Adapted protocols |  |
| --- | --- | --- | --- |
|  |  | Hypermutator<br>Ancestor | Extended<br>Subinhibitory<br>Exposure |
| Strain | PA14 | PA14 *mutS*:T114P | PA14 |
| Mutation rate (mutation/genome/division)<sup>a</sup> | 0.001 | 0.1 | 0.001 |
| Length of exposure (days)<sup>b</sup> | 3 | 3 | 10 |
| Total cell divisions<sup>c</sup> | $2.98 \times 10^9$ | $2.98 \times 10^9$ | $9.91 \times 10^9$ |
| Total mutations<sup>d</sup> | $2.98 \times 10^6$ | $2.98 \times 10^8$ | $9.91 \times 10^6$ |
| Mean mutations per nucleotide, cumulatively<sup>e</sup> | 0.458 | 45.8 | 1.52 |
| Cumulative probability of a mutation per nucleotide<sup>f</sup> | 36.8% | 100% | 73.6% |
| **Results** |  |  |  |
| Populations that survived entire experiment (%) | 0% | 50% | 40% |

### Increasing mutation supply by using a hypermutator strain promotes evolution of resistance

One way to increase the mutation supply during evolution experiments is to use an ancestral strain with a higher mutation rate. These mutator genotypes also commonly evolve during chronic infections of *P. aeruginosa* and are therefore relevant to AMR evolution (Oliver et al. 2000; Mena et al. 2008). We selected a strain of *P. aeruginosa* PA14 with a defect in mismatch repair (*mutS* T112P), which has a mutation rate ∼100 times higher than the ancestor (Flynn et al. 2016; Harris et al. 2021) and a starting level of resistance to WLBU2 of 5.3 ± 2.3 mg/L (Figure 2A). With this hypermutator strain, under the same experimental conditions described above and summing all individual cell divisions, each nucleotide experiences approximately >40 mutations over the first three days of growth under subinhibitory conditions (Table 1). Five mutator populations were propagated in each lifestyle combination following the previously described protocol. Two planktonic and three biofilm populations survived this treatment, resulting in 3 - 7.5-fold increases in resistance level relative to the ancestor (Figure 2A). The other 5 populations did not survive when exposed to the inhibitory concentration the MIC of WLBU2.

**Figure 2.**
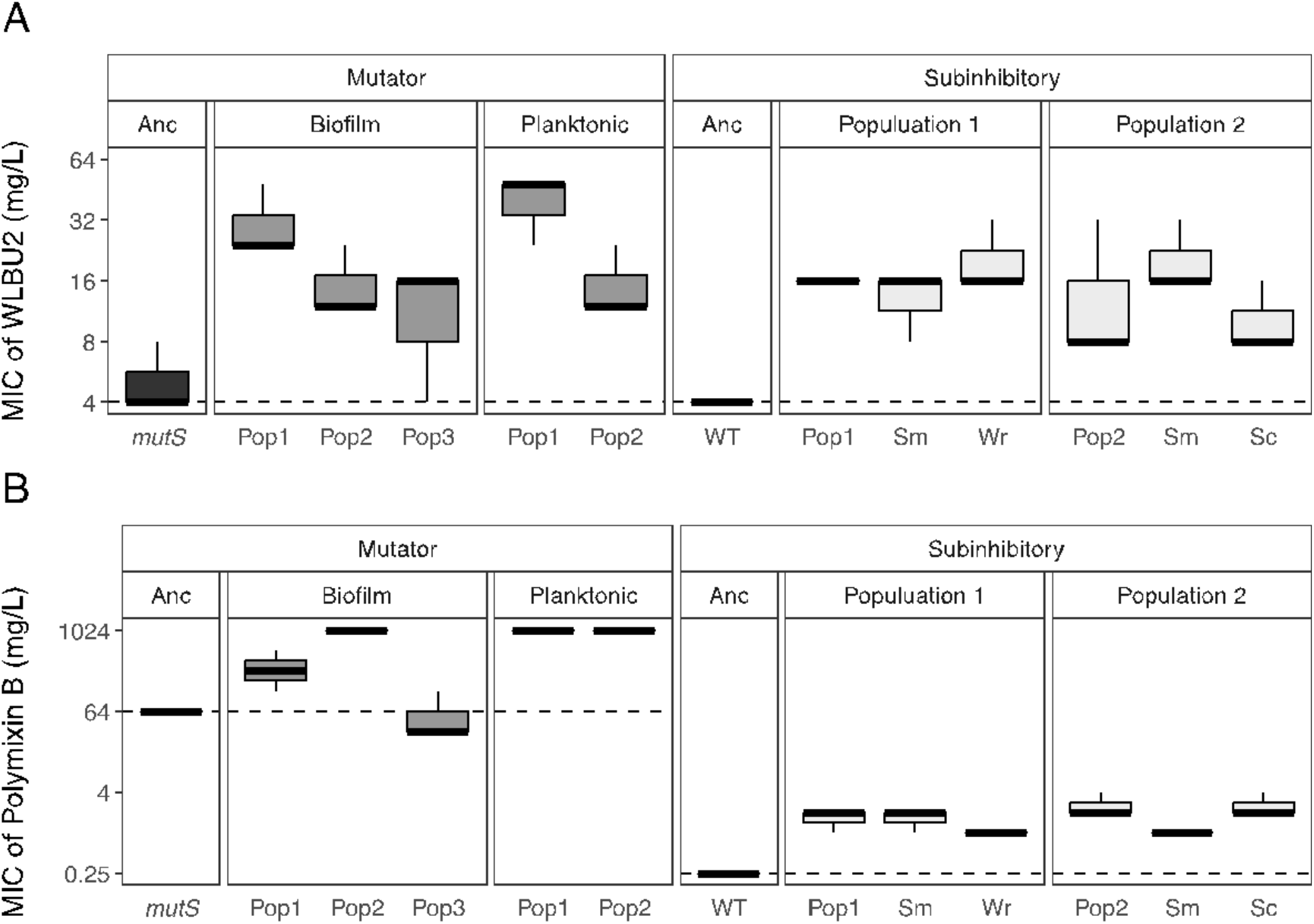
Resistance levels to WLBU2 and polymyxin B of populations evolved in the presence of WLBU2, as minimum inhibitory concentration (MIC) (mg/L). Boxplots show the median and quartiles for 3 replicates. Populations evolved from the *mutS* mutator genotype are in dark grey and resistant clones isolated from the ancestor are in light grey. All populations significantly exceed ancestral resistance to WLBU2 (p < 0.05, one-way ANOVA following Benjamini et al.) and all populations except biofilm population 3 evolved from the *mutS* ancestor significantly exceed WT resistance to polymyxin B (p < 0.05, one-way ANOVA). Dotted line denotes the MIC of the ancestor, and grey lines note maximum WLBU2 populations were challenged with during the evolution experiment.

We sequenced genomes from all five surviving populations to a depth of 148.0 ± 42.80 and detected a total of 98 mutations at frequencies > 0.1, including 31 fixed mutations (Table S3). As the ancestor was a mutator strain, it can be difficult to infer what mutations were the drivers of the resistance phenotype and which were hitchhikers (Shaver et al. 2002). We focused on instances of gene-level parallel (repeated) evolution as strong candidates, because mutations in the same gene found in independently derived lineages provide strong evidence of selection on this trait. Further, the large population size and high mutation supply empowers selection to enrich the most beneficial genotypes (Lieberman et al. 2011; Toprak et al. 2011; Vogwill et al. 2014; Bailey et al. 2015; Ibacache-Quiroga et al. 2018; Papkou et al. 2020; Scribner et al. 2020).

Only four genes, *orfN, pmrB, wspF*, and *morA*, were mutated in more than one population exposed to WLBU2 and not in the populations evolving in the absence of antibiotics, indicating roles for these mutations in evading the antibiotic treatment (Tables 2 and S3). The two-component regulator *pmrB* governs several modifications of lipopolysaccharides (LPS) and has previously been demonstrated to confer resistance to other cationic peptides (Moskowitz et al. 2012; Fernández et al. 2013). Four of the five WLBU2-resistant populations also show cross-resistance to polymyxin B (Figure 2B) increasing their MIC from 2 to ≥4 folds. The *wspF* gene encodes a methylesterase that regulates activity of the surface-sensing Wsp cluster that in turn activates the diguanylate cyclase WspR and biofilm production (Hickman et al. 2005; Huangyutitham et al. 2013). The *morA* gene encodes multiple sensor domains that control diguanylate cyclase and phosphodiesterase domains acting on the second messenger cyclic di-GMP. This molecule promotes biofilm production at high levels and motility at low levels (Hickman et al. 2005; Ha and O’Toole 2015). These biofilm-associated mutations in *wspF* and *morA* strongly indicate that production of aggregates or biofilm plays a role in resisting WLBU2.

### Extending the exposure to subinhibitory concentrations

The mutator genotype of *P. aeruginosa* facilitated the evolution of resistance to WLBU2 (Figure 2A), but isolating clones without other background mutations was impossible because of their increased mutation rate. Nonetheless, the parallelism in *orfN, morA, wspF* and *pmrB* (Figure 3) provides a strong indicator of the fitness benefits of these mutations (Lieberman et al. 2011; Vogwill et al. 2014; Cooper 2018). Because resistance could evolve by increasing the mutation supply, we tested whether increasing the number of generations in subinhibitory concentrations of WLBU2 would increase the chance for the WT (non-mutator) ancestor of acquiring mutations needed to survive inhibitory concentrations.

**Figure 3.**
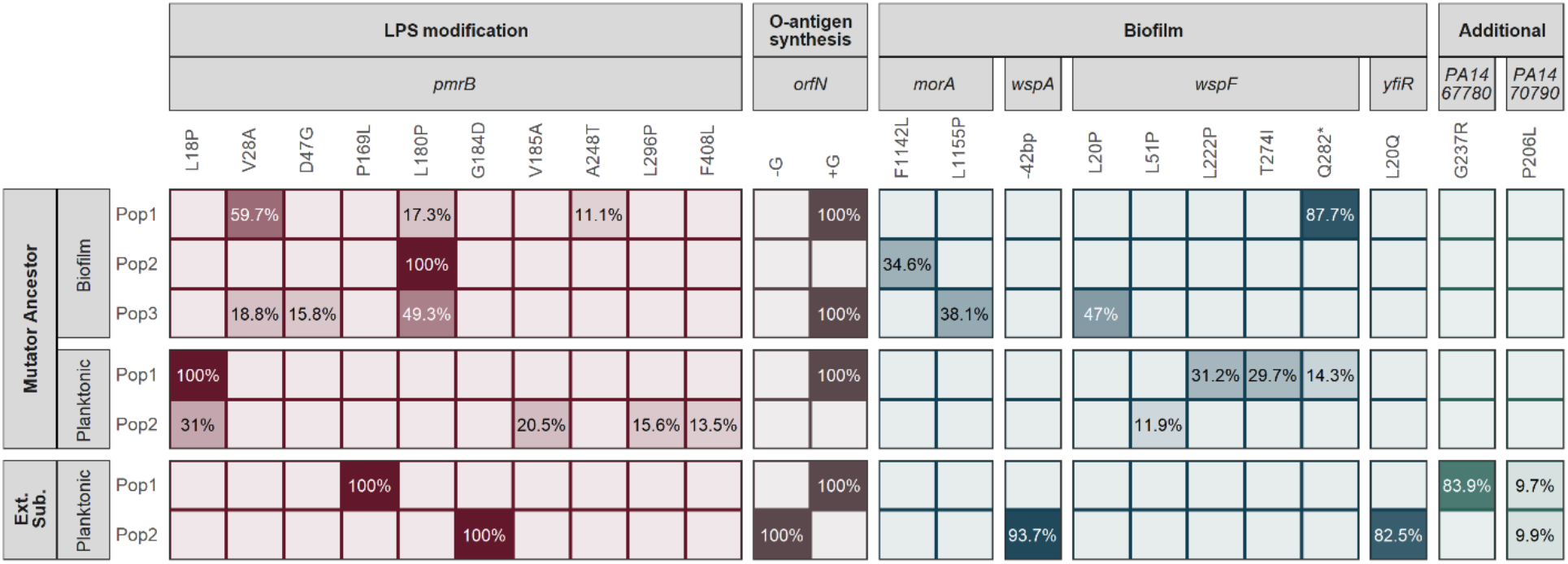
Whole genome sequencing of the populations reveal mutations in a few genes selected in parallel to high frequencies in all populations, suggesting that a minimum of two mutations among 3 key functional categories is required to increase resistance to WLBU2. While most fixed mutations were seen in the mutator-derived populations, genetic hitchhiking is more likely in these conditions. We therefore focused on parallel, high-frequency mutations seen in both the extended subinhibitory phase and mutator-founded experiments (indicative of adaptive targets and driver mutation). A complete list of mutations is shown in Tables S3 and S4. More details of parallel mutations are shown in Table S5.

We propagated five planktonic populations of WT PA14 (MIC to WLBU2 = 4 ± 0 mg/L, Figure 2) for 10 days under subinhibitory concentrations of WLBU2, followed by 2 days at inhibitory concentrations (Figure 1). We estimate that the probability that any given nucleotide would be mutated during this regime is 0.74, with mean mutations per site of 1.52 (Table 1). Again, as a control, three populations were propagated in the absence of WLBU2 to distinguish between mutations adaptive to broth or WLBU2. Only two populations survived the prolonged subinhibitory treatment. Those two populations showed 3-4-fold increased resistance to WLBU2 and 2-3-fold increases to polymyxin B (Figure 2). We also detected different colony morphologies within each population at the end of the experiment. Population 1 included small, rugose or wrinkly (Wr) colonies in addition to the smooth (Sm) morphology of the ancestor, and population 2 contained both Sm and Wr colonies (Figure 4). It is interesting to note that small colony variants are associated with aggregation, increased biofilm formation, and worse outcomes in chronic infections (Fux et al. 2005; Gloag et al. 2019). We tested whether these clones produced more biofilm by measuring aggregation in the presence or absence of WLBU2. We found that the Wr clones settled in clumps more than the ancestral strain both in absence and presence of WLBU2, while Sm colonies only clustered more in the presence of the peptide (Figure 5).

**Figure 4.**
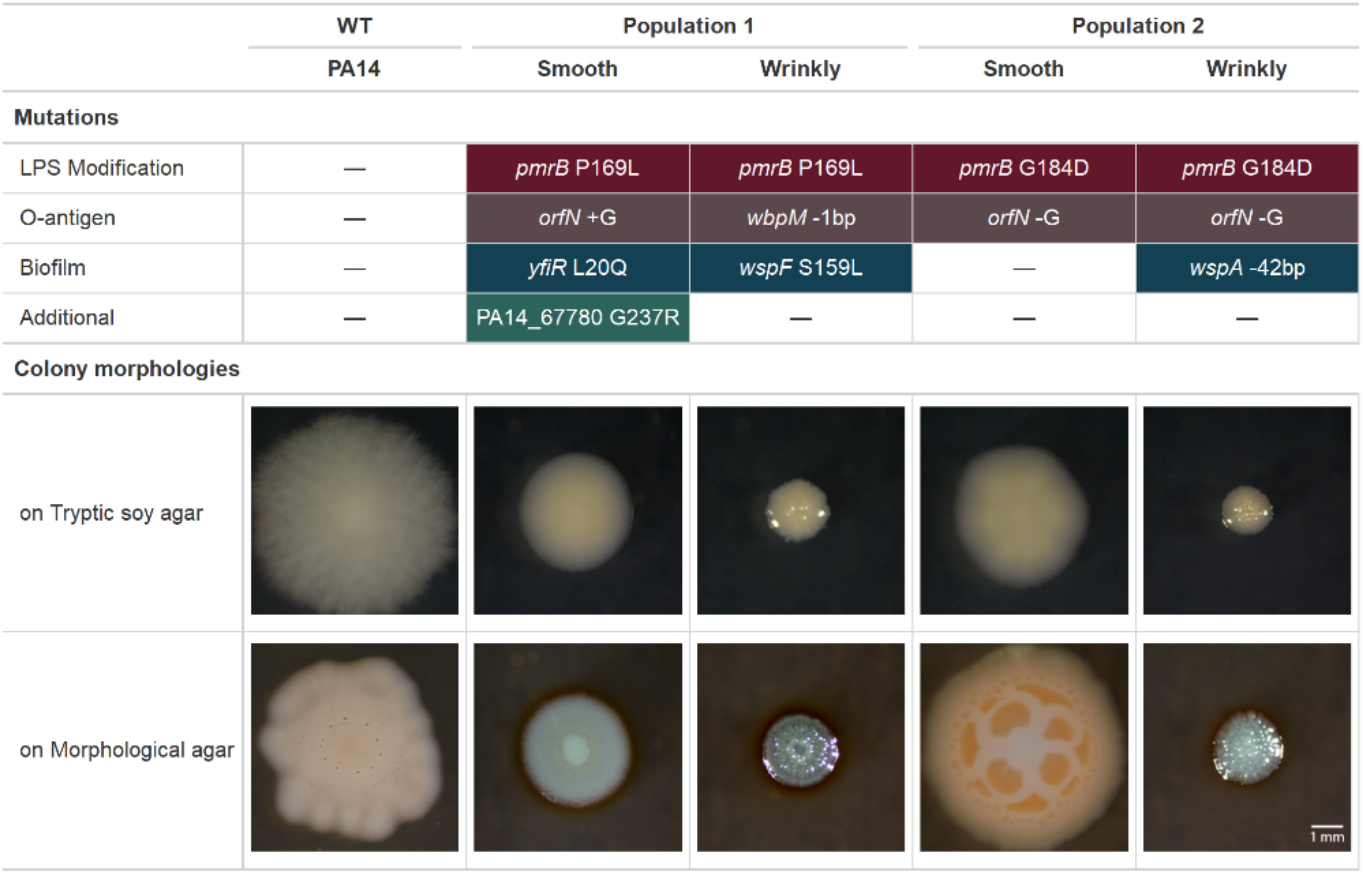
Isolated clones have at least two mutations involving LPS modification and O-antigen synthesis, some with additional biofilm mutations, affecting colony morphologies. Each wrinkly clone has a specific mutation that was not detected at the population level (*wbpM* and *wspA*), but they still fall into the functional categories associated with resistance shown in Fig 3. The two wrinkly colonies have expected *wsp* mutations, while the two smooth morphotypes differ from their wild-type ancestor in size, shape, and coloration. It is notable that 3 of 4 isolates acquired biofilm-associated mutations despite no explicit biofilm selection.

**Figure 5.**
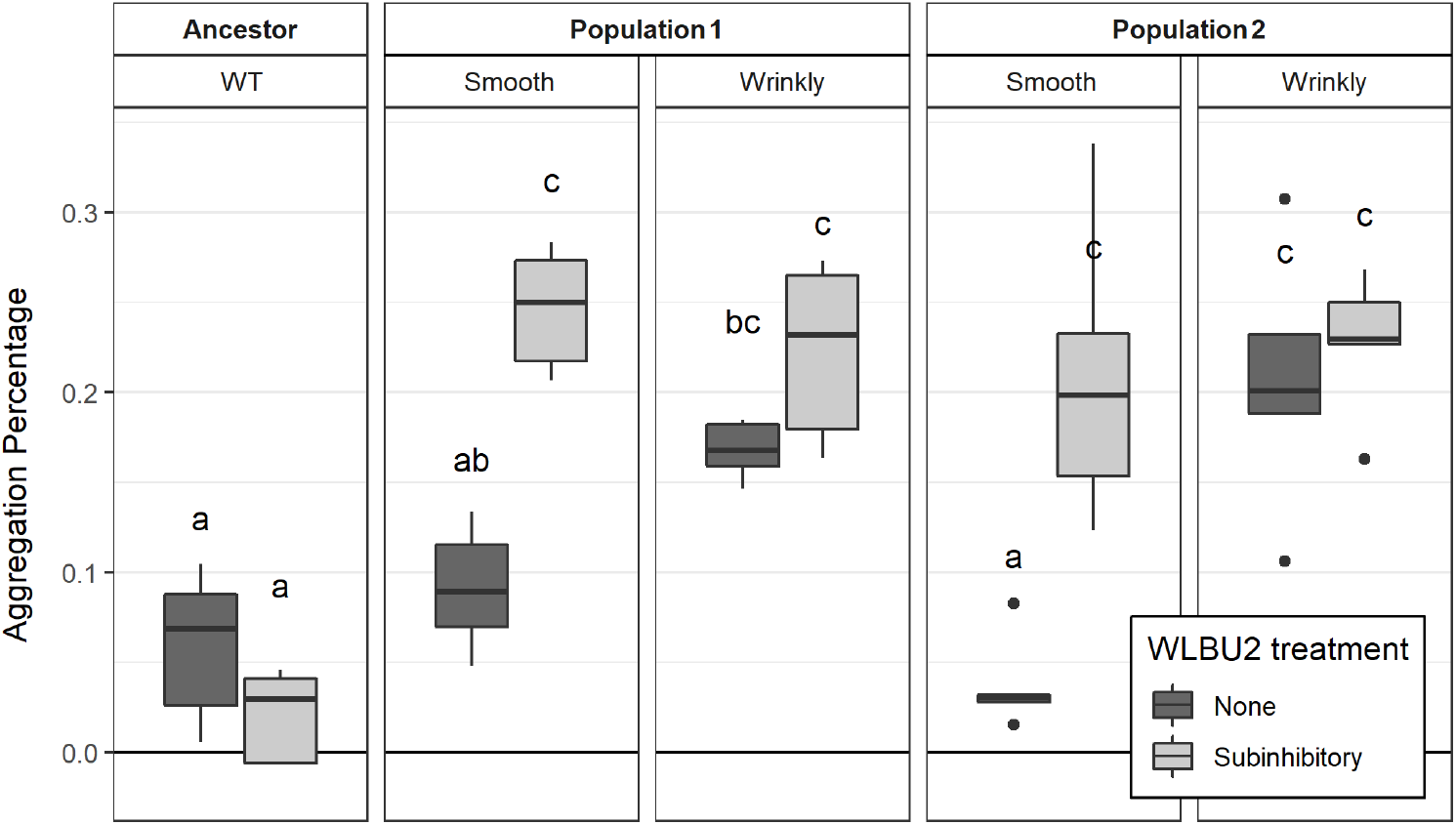
Aggregation of PA14 and resistant clones. The resistant Wrinkly (Wr) clones (associated with *wsp* mutations) aggregate more than WT regardless of WLBU2 treatement. The Smooth (Sm) clones did not differ from WT in the absence of WLBU2, but aggregated when exposed to WBLU2. Aggregation was measured as settling for 24 hours at 4°C, n=5. Means sharing a letter are not significantly different (Sidak-adjusted least-square means, p <0.05). Dark and light grey denote culturing in the absence and presence of WLBU2 respectively.

WGS of the two surviving populations and representative clones of each colony type revealed that all acquired mutations in *pmrB* as well as mutations known to increase biofilm production (Figure 3). The two small colony variants acquired mutations in *wspA* or *wspF* in the Wsp pathway seen previously in the mutator experiment, while one resistant Sm clone acquired *orfN* and *pmrB* mutations as well as *yfiR*, another gene known to activate c-di-GMP synthesis (Li et al. 2015). It is notable that these biofilm-activating mutations were selected during planktonic propagation of the populations, suggesting that forming aggregates is a key step for resistance to WLBU2. In addition, both resistant populations and the two clones with smooth morphology had mutations in the *orfN* gene, which is involved in the synthesis of the O-antigen (Rocchetta et al. 1999). Further, one Wr isolate had a mutation in *wbpM*, which also contributes to O-antigen biosynthesis (Rocchetta et al. 1999).

## Discussion

The rapid emergence and dissemination of encoded resistance mechanisms is decreasing the effectiveness of antibiotics in the clinic. Antimicrobial peptides (AMPs) are components of the innate immune systems of many animals and plants and have been proposed as good candidates for treating multidrug resistant infections where other antibiotics have failed (Deslouches et al. 2015; Mahlapuu et al. 2016; Lei et al. 2019). AMPs derived from natural defensive peptides have advantages compared to traditional antibiotics thanks to their broad-spectrum activity, ability to treat biofilm-embedded populations and observed evolutionary constraints that limit the rapid acquisition of AMP resistance (Lei et al. 2019; Deslouches et al. 2020; Jangir et al. 2021). In fact, numerous AMPs, including daptomycin, vancomycin and bacitracin, have been approved by the FDA, and several other AMPs, including WLBU2, are under clinical trials (Greber and Dawgul 2017). However, AMPs face some handicaps that have limited their successful development for clinical use, including low activity in acidic conditions such as blood or plasma, host toxicity, susceptibility to protease digestion and poor absorption (Falagas and Kasiakou 2005; Deslouches et al. 2020). Nevertheless, recent research with several engineered AMPs have demonstrated that most of these limitations can be overcome by structural re-design (Cirac et al. 2017; Rabanal and Cajal 2017; de Breij et al. 2018; Mwangi et al. 2019; Deslouches et al. 2020; Torres et al. 2021). WLBU2 is an engineered AMP derived from the natural LL-37 peptide that has overcome most of the limitations described above (reviewed in (Deslouches et al. 2020)). The effectiveness of WLBU2 against a range of bacterial species in multiple conditions and the absence of clear genetic mechanisms for resistance makes it a promising candidate as a new antibiotic. However, one exception is its relative ineffectiveness against the *Burkholderia cepacia* complex (Deslouches et al. 2015), which is generally resistant to the cationic peptide polymyxin B. Despite this exception, the lack of any defined resistance mechanism suggests that multiple modifications to LPS or outer envelope (e.g. associated with major species differences rather than variation at a single locus) may be required for WLBU2 resistance. If multiple mutations are indeed necessary for WLBU2 resistance, this should substantially lower the probability of evolved resistance during treatment. This study provides strong support for this multiple mutation model, but also shows that the pathway to resistance is plausible for mutator strains or ones already containing one or more contributing mutations. Either possibility has been demonstrated in *P. aeruginosa* infections: mutator isolates are frequently reported (Oliver et al. 2000; Mena et al. 2008), and mutations in the primary driver gene *pmrB*, as well as those in the Wsp pathway or affecting O-lipid biosynthesis are also common adaptations in clinical isolates. These mutations can be selected for their advantages in biofilms even in the absence of antibiotics (Starkey et al. 2009; Moskowitz et al. 2012; Cannatelli et al. 2017; Gloag et al. 2019).

By exposing bacterial populations to antibiotics in controlled environments we can understand the routes that lead to resistance (Martínez et al. 2011). When new mutations are found in parallel in independently propagated lines in different strains and under conditions (for instance biofilm or planktonic) and not in the controls, then those mutations are the cause of the new heritable resistant phenotype (Barrick and Lenski 2013). After sequencing all populations that survived the WLBU2 treatments (5 using the mutator lineages, and 2 propagated under prolonged subinhibitory selection), the only mutations common across lineages occurred in *pmrB, orfN, morA* and *wsp* (Table S5), providing clear evidence of fitness benefits of these mutations affecting both LPS composition and biofilm production in the presence of WLBU2 and their implication in the increase of resistance to WLBU2.

It has been well demonstrated that resistance to cationic peptides in *P. aeruginosa* involves modification of LPS (Fernández et al. 2013). Consistent with these findings, we detected parallel evolution of mutations in the *pmrB* and *orfN* genes across four independent lineages. PmrB forms part of a two-component regulatory system that modifies the lipid-A composition including its negative charge, a mechanism that is commonly associated with cationic peptide resistance in several species (Moskowitz et al. 2012; Cannatelli et al. 2017). In combination with *pmrB* variants, mutations in *orfN* and *wbpM* were found in WBLU2-resistant populations and clones. OrfN and WbpM form part of the operon that synthetizes the LPS O-antigen, and mutations in *orfN* are predicted to increase antibiotic resistance by reducing membrane permeability (Rocchetta et al. 1999; Scribner et al. 2020). Furthermore, we observed mutations in *wspA, wspF*, and *yfiR* that increase aggregation and/or biofilm production, likely through increased cyclic-di-GMP that in turn increases production of the cationic Pel polysaccharide. It will be important to explore how this positively charged component of the biofilm matrix interacts with the positively charged WLBU2 peptide to provide defense. Mutations in the phosphodiesterase domain of *morA* also were repeatedly selected in mutator populations with a probable similar effect on cyclic-di-GMP and Pel-mediated biofilm production (Katharios-Lanwermeyer et al. 2021). Aggregation by secretion of a biofilm polymer is therefore another likely mechanism of WLBU2 resistance or tolerance (Starkey et al. 2009; Jennings et al. 2015; Gloag et al. 2019).

As AMR spreads around the globe, it is crucial to develop new antibiotics so that we still have treatments available for otherwise routine infections. During antibiotic development, we must anticipate possible mechanisms of resistance in their design (Brockhurst et al. 2019; MacLean 2020). Furthermore, to broaden use of AMPs in clinical settings we must explore evolutionary, genetic, and phenotypic barriers to AMP resistance in relevant pathogens (Jangir et al. 2021). We often read claims of new ‘evolution-proof’ antibiotics (Zasloff 2002; Ling et al. 2015), but here we further demonstrate that the absence of resistance mechanisms can result from insufficient sampling of genetic diversity (Bell and MacLean 2018). We also reveal the genetic causes of evolved resistance to the promising new cationic peptide WLBU2 by combining experimental evolution under multiple population-genetic conditions with genome sequencing of whole populations and resistant clones. Our findings indicate that WLBU2 likely has the advantageous property of requiring two or more mutations that affect the charge of the outer membrane as well as cellular aggregation for resistance to evolve. Further biochemical assays are needed to determine the discrete roles of each mutation in the resistance phenotype. This additional research notwithstanding, we believe that effort should be spent elucidating the evolutionary pathways driving antibiotic resistance when developing antibiotics, because knowing the likely adaptations could help in designing more accurate treatments, limit the spread of AMR, and preserve the longevity of the antibiotic in clinical use.

## Methods

### Experimental evolution

We performed three independent evolution experiments. In all of them, we propagated the ancestor strains in a rich medium modified from M9 (referred to as M9^+^) containing 0.37 mM CaCl_2_, 8.7 mM MgSO_4_, 42.2 mM Na_2_HPO_4_, 22 mM KH2PO_4_, 21.7mM NaCl, 18.7 mM NH_4_Cl and 0.2 g/L glucose and supplemented with 20 mL/L MEM essential amino acids (Gibco 11130051), 10 mL/L MEM nonessential amino acids (Gibco 11140050), and 10 mL each of trace mineral solutions A, B, and C (Corning 25021-3Cl).

In the first experiment, we selected a single clone of *A. baumannii* ATCC17978 and *P. aeruginosa* PA14 and propagated them independently for 24 hours in M9^+^ in the absence of antibiotic. We then sub-cultured each population into 10 replicate populations. Ten of the populations (5 planktonic and 5 biofilm) were propagated every 24 hours in increasing concentrations of WLBU2 starting at 0.5X the minimum inhibitory concentration (MIC). We doubled the WLBU2 concentrations after 72 hours until no population survived the treatment. As a control, we propagated the same number of strains both planktonically and in biofilm in the absence of antibiotic. We also propagated 10 populations (5 biofilm and 5 planktonic) in increasing concentration of the cationic peptide polymyxin B.

In the second experiment, we propagated 5 planktonic populations and 5 biofilm populations under the same conditions described above, a mutator strain of *P. aeruginosa* obtained from (Flynn et al. 2016). We also propagated 3 planktonic populations and 3 biofilm populations of the hypermutant strain in the absence of antibiotic pressure.

In the third experiment, we propagated 5 planktonic populations of the non-hypermutant laboratorial strain *P. aeruginosa* PA14 under subinhibitory concentrations of WLBU2 for 10 days. Then, we propagated the populations that survived the treatment with 1X MIC of WLBU2 2 days.

All the experimental evolutions were performed using 18 mm glass tubes (Figure S1). For the planktonic propagation, we serial passaged 50 µl into 5 ml of M9^+^ (dilution factor 100) which corresponds to approximately 6.64 generations per day (Lenski 1991). For biofilm populations, we transferred a polystyrene bead (Polysciences, Inc, Warrington, PA) to fresh media containing three sterile beads. We rinsed each bead in PBS before the transfer, therefore reducing the transfer of planktonic cells. Each day we alternated between black and white marked beads, ensuring that the bacteria were growing on the bead for 24 hr, which corresponds to approximately 6 to 7.5 generations/day (Turner et al. 2018).

In all three experiments, we froze 1mL of the surviving populations at days 1, 3, 4, 6, 7, 9, 10, and 12 in 9% of DMSO (before and after increases in antibiotics.)

### Phenotypic characterization: antimicrobial susceptibility and aggregation assay

We determined the MIC to WLBU2 and polymyxin B of the whole populations and 4 clones by broth microdilution according to the Clinical and Laboratory Standards Institute guidelines (CLSI 2019), in which each bacterial sample was tested in 2-fold-increasing concentrations of each antibiotic. WLBU2 was provided by Peptilogics (Peptilogics, Pittsburgh, PA) and polymyxin B was provided by Alfa Aesar (Alfa Aesar, Wardhill, MA). We performed this experiment with three replicates in the of antibiotic.

To determine the aggregation ability of the 4 clones selected in the subinhibitory experiment, we grew two replicates (R1 and R2) of each clone and the ancestral strain in 5ml of M9+ for 24 hours at 37°C in a roller drum at 200 RPM. After 24 hours we transferred the whole culture of each clone to a 13 mm glass tube, and let the tubes settle over 24 hours at 4°C without shaking. We vortexed replicate 1 (R1) of each clone for 30s and measured the OD_600_. From R2, we carefully took 200 μl of the upper fraction without vortexing and measured the OD_600_. The aggregation percentage was estimated as 100(1 - OD_600_ R2 / 1 – OD_600_ R1). We performed this experiment with five replicates both in the absence and presence (4 μg/ml) of WLBU2.

### Genome sequencing

We performed whole population genome sequencing of population that survived the treatment in each experiment as well as control lines with a coverage of 148.01 ± 42.80. This includes 3 biofilm and 2 planktonic populations of mutator populations surviving WLBU2 treatment and 4 biofilm and 3 planktonic populations evolved in the absence of the antibiotic (12 total populations). In addition, we sequenced the 2 populations and 4 clones that survived the subinhibitory WLBU2 treatment, as well as one untreated control population (2 populations, 4 clones). Finally, we sequenced 3 biofilm and 3 planktonic populations of *A. baumannii* that were propagated in presence of polymyxin B as well 3 three planktonic and 3 biofilm populations (12 total populations).

We revived each population or clone from a freezer stock in the growth conditions under which they were isolated and grew for 24 hours. DNA was extracted using the Qiagen DNAeasy Blood and Tissue kit (Qiagen, Hiden, Germany). The sequencing library was prepared as described by Turner and colleagues (Turner et al. 2018) according to the protocol of Baym *et al*. (Baym et al. 2015), using the Illumina Nextera kit (Illumina Inc., San Diego, CA) and sequenced using an Illumina NextSeq500.

### Data processing

The variants were called using the breseq software package v0.31.0 (Barrick et al. 2014) using the default parameters and the -p flag when required for identifying polymorphisms in populations after all sequences were first quality filtered and trimmed with the Trimmomatic software v0.36 (Bolger et al. 2014) using the criteria: LEADING:20 TRAILING:20 SLIDINGWINDOW:4:20 MINLEN:70. The version of *A. baumannii* ATCC 17978-mff (GCF_001077675.1 downloaded from the NCBI RefSeq database,17-Mar-2017) was used as the reference genome for variant calling. We added the two additional plasmid sequences present in the *A. baumannii* strain (NC009083, NC_009084) to the chromosome NZ_CP012004 and plasmid NZ_CP012005. The version of *P. aeruginosa* UCBPP-PA14 was downloaded from RefSeq on 25-Aug-2020. To remove false positives using the following strategy: mutations were removed if they never reached a cumulative frequency across time points of 25%, or if mutations were also found in the ancestor’s genome. The *mutS* clone used in this study is a merodiploid bearing both the ancestral and the mutated *mutS* gene (Flynn et al. 2016). As breseq can fail to detect mutations when analyzing repeated sequences (Barrick et al. 2014), we manually visualized the *mutS* locus with the bam files generated by breseq using the Integrative genomics viewer software, confirming that the mutation was present in the ancestral clone and in all evolved populations sequenced at *ca*. 50% frequency (Robinson et al. 2011).

### Data sharing statement

R code for filtering and data processing are deposited here: https://github.com/sirmicrobe/U01_allele_freq_code/tree/master/wlbu2_paper. All sequences were deposited into NCBI under the Bioproject accession number PRJNA663022.

## Acknowledgments and funding information

We thank Jonathan Steckbeck, Ken Urish and Peptilogics for kindly providing WLBU2 and for their technical assistance. We thank Ken Urish, Jeronimo Rodriguez-Beltran, Alecia Rokes and Cristina Herencias for proofreading of the paper. This research was supported by NIH U01AI124302-01 and, in part, under PA CURES Grant #4100085725 with the Pennsylvania Department of Health. The Department specifically disclaims responsibility for any analyses, interpretations or conclusions. AS-L is supported by the European Commission (H2020-MSCA-IF-2019, 895671-REPLAY). VAH and AHPB were supported by the National Institute of General Medical Sciences of the National Institutes of Health under Award Number T32GM008208. The content is solely the responsibility of the authors and does not necessarily represent the official views of the National Institutes of Health.

## Transparency declaration

None to declare.

---

**Table S1**. Mutations selected in the presence of increasing concentrations of polymyxin B. Two sheets can be found in the Table S1: “PolymyxinB_raw” which includes all mutations called by breseq and “PolymyxinB_relevant” which include the mutations selected for in presence of polymyxin B but not selected in the absence of the antibiotic.

**Table S2**. Estimated mutation probabilities during experimental evolution.

**Table S3**. Mutations selected using the *mutS* deficient strain. Three sheets can be found in the document: “raw_mutations” include all mutations called by breseq, “no_controls_>0.1” include the list of mutations subtracting from the raw list all mutations happening in the populations evolving without WLBU2 and with lower frequency than 0.1, and “parallel_genes” which include the three genes that are targeted in more than one resistant population and are not present in any control.

**Table S4**. Mutations selected in the subinhibitory experiment. Two sheets can be found in the document: “populations” and “clones” include all mutations called by breseq in the evolving populations and clones respectively. Note that the curated list of mutations, subtracting those found in controls without antibiotic and the ancestral strain, is listed in Figure 2.

**Table S5**. Instances of parallel evolution events across the experiments. Four sheets are automated filtering and grouping of the raw mutations called by breseq for both mutS-founded and extended subinhibitory experiments (with mutations common to the ancestor removed) to highlight instances of parallelism: “All_mutations_>0.1” is all site-specific mutations at 10% or greater in any population, “All_mutations_>0.1_bygene” is this same list grouped by gene with cumulative frequencies, “Parallel_grouped_by_site” lists all site-specific mutations called in more than 1 population, and “parallel_grouped_by_gene” lists genes targeted across multiple mutations. “Grouped_bySite_Cleaned” lists the site-specific mutations seen in multiple populations and those seen at greater than 50% in the extended subinhibitory experiment, with erroneous mutation calls due to mismappings manually removed. “Grouped_byGene_Cleaned” further generalizes this curated list to the gene level with cumulative frequencies.

